# Robust and accurate decoding of hand kinematics from entire spiking activity using deep learning

**DOI:** 10.1101/2020.05.07.083063

**Authors:** Nur Ahmadi, Timothy G. Constandinou, Christos-Savvas Bouganis

## Abstract

Robustness and decoding accuracy remain major challenges in the clinical translation of intracortical brain-machine interface (BMI) systems. In this work, we show that a signal/decoder co-design methodology (exploiting the synergism between the input signal and decoding algorithm within the design development process) can be used to yield robust and accurate BMI decoding performance. Specifically, through applying this process, we propose the combination of using entire spiking activity (ESA) as the input signal and quasi-recurrent neural network (QRNN) based deep learning as the decoding algorithm. We evaluated the performance of ESA-driven QRNN decoder for decoding hand kinematics from neural signals chronically recorded from the primary motor cortex area of a non-human primate. Our proposed method yielded consistently higher decoding performance than any other methods previously reported across long-term recording sessions. Its high decoding performance could sustain, even when spikes were removed from the raw signals. Overall results demonstrate exceptionally high decoding accuracy and chronic robustness, which is highly desirable given it is an unresolved challenge in BMIs.

## Introduction

Millions of individuals worldwide suffer from either partial or complete paralysis due to neurological disorders such as spinal cord injury (SCI), amyotrophic lateral sclerosis (ALS), and stroke. For paralysed individuals, regaining arm and hand movements is one of the most important outcomes of any clinical treatments.^1,2^ Intracortical brain-machine interfaces (BMIs) have emerged as a promising clinical tool for restoring lost motor functions by translating neural activity directly into control signals for guiding assistive devices.^3,4^ Early pilot clinical studies have demonstrated that BMIs can be used to control a computer cursor,^5–8^ robotic arm,^9–11^ or functional electrical stimulation (FES) system.^12–14^ While there has been tremendous progress in the field of BMIs from both nonhuman primate and human studies in the past two decades, several challenges remain to be overcome to achieve a clinically viable translation of BMIs. Among them are the robustness and accuracy of BMI systems,^15–18^ which are greatly influenced by the type of input signal and decoding algorithm (i.e. decoder) being used.

BMIs have predominantly utilised spikes or action potentials “fired” by individual neurons which are also known as single-unit activity (SUA) as the input signal.^19–21^ However, numerous studies have shown that spike recordings are not chronically stable, with measured signals fading, and number of observable units decreasing over time,^22–24^ which in turn leads to degraded performance. The instability and diminishing trend of recorded spikes are thought to be primarily caused by the foreign body response inducing glial scarring around the electrodes, micromotion of the electrodes, and degradation of insulation properties.^25–28^ One approach to overcome this issue is by utilising a different type of input signals, namely multiunit activity (MUA). MUA, defined as all spikes detected through threshold crossing without spike sorting, offers simpler processing while providing better signal stability over time than SUA.^25, 29, 30^ Another alternative input signal is local field potential (LFP), which is thought to mainly reflect summed synaptic activity from a local neuronal population around the recording electrodes. It is believed and has been demonstrated by experimental studies that LFPs exhibit better long-term signal stability than their spike counterparts.^31–34^ Moreover, LFPs can be obtained by simpler processing and lower sampling rate that can reduce the power consumption of BMIs. Despite the appealing advantages of MUA and LFP, a considerable number of published studies have reported that the decoding accuracy of MUA^24, 29, 30, 35, 36^ and LFP^33, 36–39^ are lower than that of SUA. Therefore, it is highly desirable to have an input signal that is not only stable but also yields high decoding accuracy.

The decoding accuracy is also affected by the decoder, that is, an algorithm used to convert the input signal from the brain into the behavioural parameter of interest (e.g. hand kinematics). Many BMI studies employ linear decoders, e.g. Wiener filter (WF)^5, 40, 41^ and Kalman filter (KF),^6, 9, 21, 32, 33, 42^ which could yield suboptimal decoding accuracy as neural signals are known to exhibit nonlinear and nonstationary properties.^43^ Although there exists a nonlinear extension of WF and KF, called Wiener cascade filter (WCF)^19, 39, 44^ and unscented Kalman filter (UKF),^45, 46^ WCF and UKF assume that the noise is (additive) stationary and Gaussian, respectively. If these a priori assumptions are violated, the decoding performance of both decoders will not be optimal.

The rise of artificial intelligence, particularly deep learning (DL), with its state-of-the-art performance across a wide range of tasks (e.g. image and speech recognition, language translation, and medical diagnostics),^47, 48^ has presented the opportunity to potentially improve the decoding performance of BMIs. The question is how can this best be applied, and how significant an impact can this have on the overall performance. Recent studies have demonstrated that different variants of deep learning decoders, such as standard recurrent neural network (SRNN), long short-term memory (LSTM), and gated recurrent unit (GRU), outperform classical decoders,^49–52^ Despite this, the amount of BMI research utilising deep learning is still in its infancy. A parallel is perhaps that of voice recognition in the 1980s versus today. Standalone voice recognition software originally required significant training for each user, was sensitive to variations in tone or speed, and made frequent errors. This hindered it completely unusable. With the advent of cloud services and big data, platforms leveraging on DL (e.g. Amazon Alexa, Apple Siri) could provide robust recognition across multiple users, accents, varying tone, with background noise, etc. With improved standardisation in data collection and the availability of more data, DL now has the opportunity to impact neural decoding in similar ways.

Thus far, the selection of the input signal, and decoding algorithm have been addressed in isolation, when improving the robustness and accuracy of BMIs. DL here provides a true opportunity to holistically optimise the end-to-end system. This study is the first to address these both issues simultaneously by exploiting the synergism between the input signal and decoding algorithm in the design process. We refer to this approach as signal/decoder co-design methodology. Specifically, through applying this process, we simultaneously propose entire spiking activity (ESA) as the input signal and DL based on quasi-recurrent neural network (QRNN) as the decoding algorithm. We evaluated the performance of ESA-driven QRNN decoder for decoding (offline) hand kinematics from neural signals chronically recorded with an intracortical microelectrode array implanted in the primary motor cortex area of a Rhesus macaque monkey. We show that our proposed method achieved significantly higher decoding performance than combinations of other input signals (SUA, MUA, and LFP) and decoders (WF, WCF, KF, UKF, SRNN, LSTM, and GRU). We also show that the proposed method could sustain higher decoding performance over time under various conditions, such as (1) when spikes were removed from the raw signals, (2) the different number of channels or features, and (3) the smaller amount of training data.

## Results

### Characteristic comparison across neural signals

Firstly, we quantified the characteristics of stability and quality of SUA and MUA over time with three metrics: (1) number of units, (2) average peak-to-peak voltage amplitude (Vpp; in *µ*m) across units, and (3) mean spike rate (Hz) across units. Since the number of channels for LFP and ESA remained the same (96 channels) throughout the recording sessions, to quantify the characteristics of LFP and ESA, we only used their mean amplitude (*µ*m) across channels. Fig. 2 shows the characteristic comparison across different neural signals. We found statistically significant decreasing trends for SUA in all three metrics: the number of units declined by 11.19% (* *p* < 0.05), the Vpp declined by 18.26% (*** *p* < 0.001), and the mean spike rate declined by 0.11% (* *p* < 0.05) per day, as shown in Fig. 2a,b,c, respectively. As for MUA (see Fig. 2a,b,d), the number of units and Vpp increased by 1.48% and 186.57% per day, respectively, whereas the mean spike rate statistically significantly decreased by 0.56% per day (*** *p* < 0.001). In the case of LFP, the mean amplitude was found to increase by 1.27% per day (Fig. 2e). We did not observe a statistically significant decreasing trend in the mean amplitude of ESA although the rate was negative (−0.88% per day) as can be seen in Fig. 2f.

**Fig. 1.**
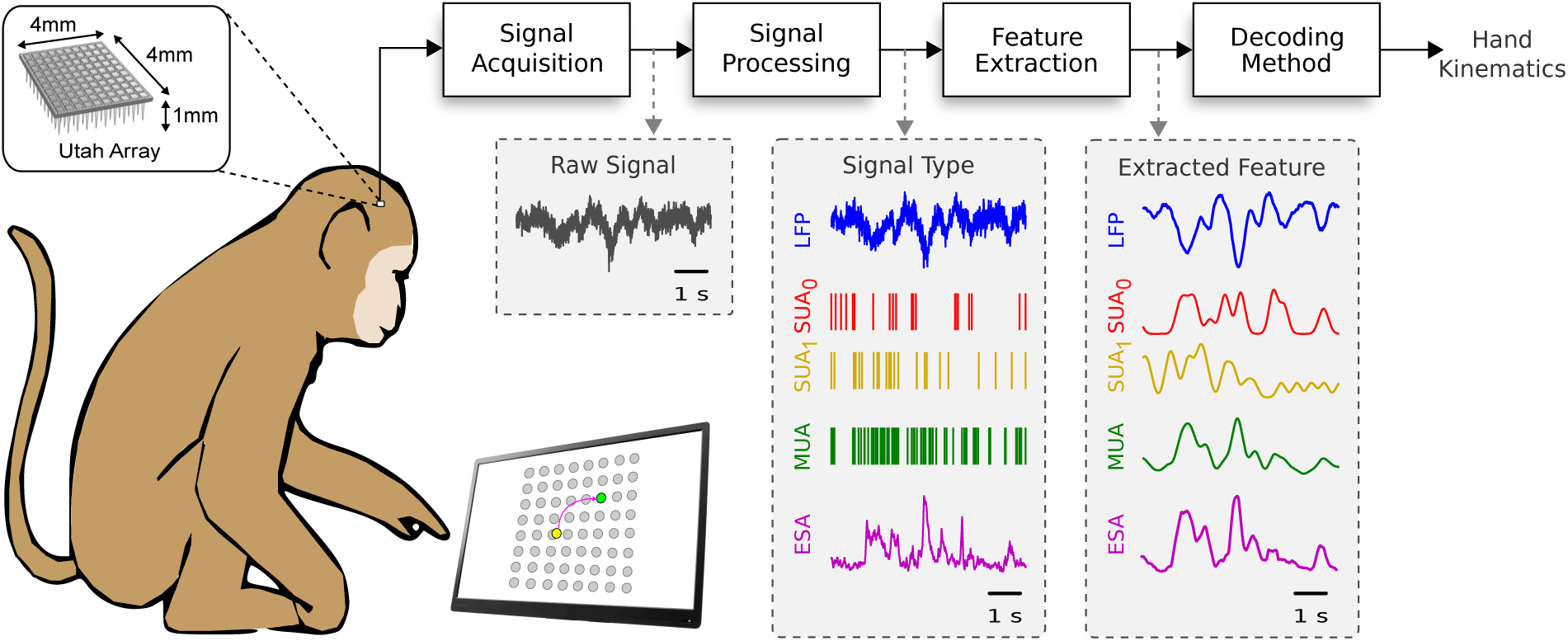
Schematic diagram of the brain-machine interface (BMI) system. Raw neural signals were recorded using a 96-channel Utah microelectrode array from primary motor cortex (M1) area of a monkey while performing continuous reaching tasks. Four different types of neural signals were obtained from the raw neural signals, namely local field potential (LFP), single-unit activity (SUA), multiunit activity (MUA), and entire spiking activity (ESA). Features extracted from these neural signals were used as inputs for different decoding methods.

**Fig. 2.**
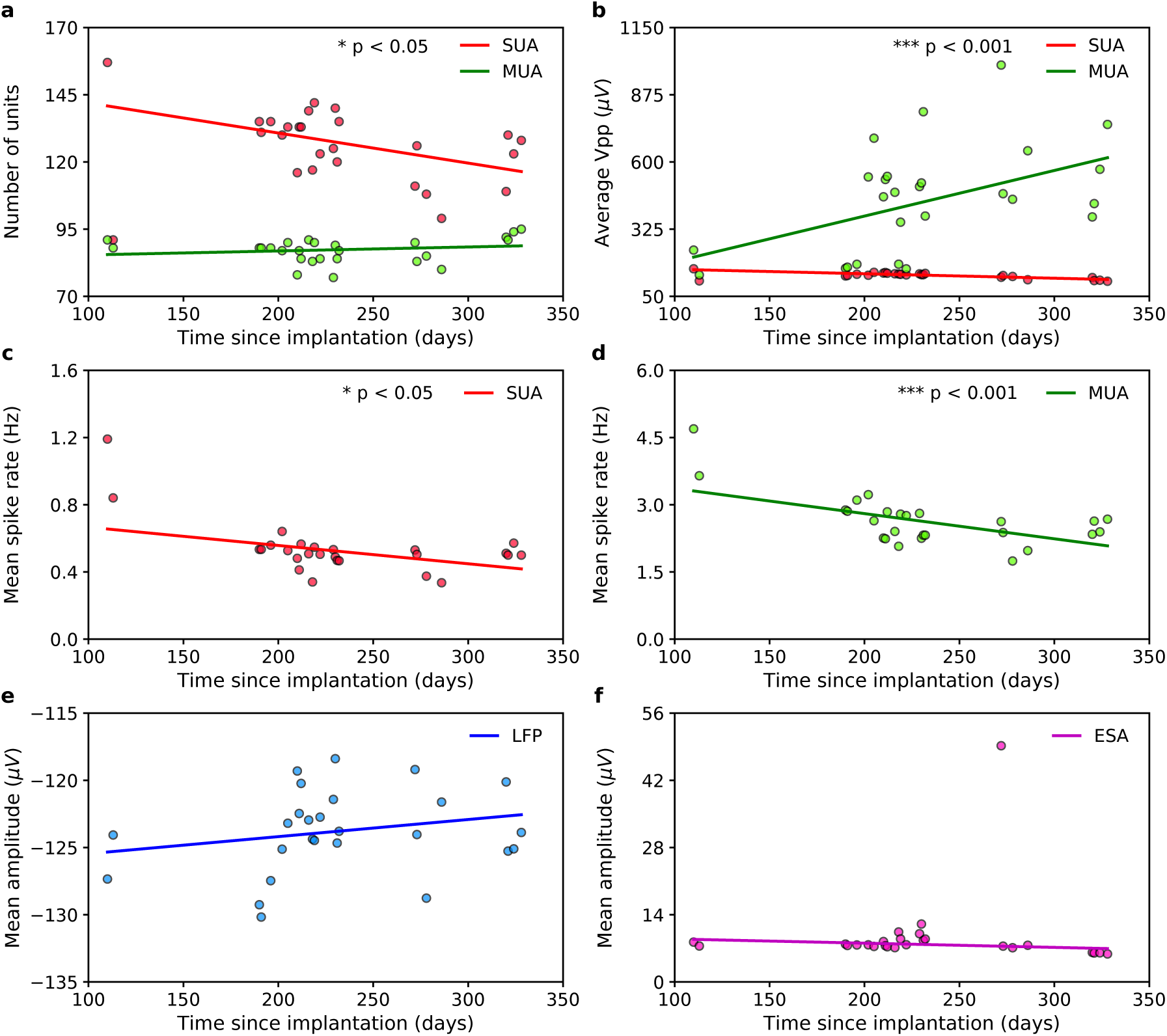
Long-term stability of different types of neural signals. **a**, Number of SUAs and MUAs over time. **b**, Average peak-to-peak voltage amplitudes (Vpp; in *µ*m) of SUA and MUA over time. **c,d**, Mean spike rates (Hz) of SUA and MUA over time, respectively. **e,f**, Mean amplitudes (*µ*m) of LFP and ESA over time, respectively. Huber based robust linear regression was used to test whether or not the trends significantly linearly decreased. Asterisks represent the trends of neural signals that significantly linearly decreases (one-tailed one-sample *t*-test; * *p* < 0.05, *** *p* < 0.001).

We then examined the redundancy characteristics of neural signals by measuring their inter-channel/unit correlation. For each neural signal, we calculated the correlation between all possible pairs of units/channels and averaged them across units/channels. The average inter-channel/unit correlation for all neural signals is displayed in Fig. 3. SUA was found to have the lowest average correlation (0.10 ± 0.01; mean ± SD), whereas LFP had the highest average correlation (0.59 ± 0.09) as can be seen in Fig. 3a and Fig. 3d, respectively. On the other hand, MUA and ESA had an average correlation of 0.25 ± 0.05 and 0.45 ± 0.12, respectively, as shown in Fig. 3b and Fig. 3c, respectively. By sorting according to the average correlation scores, we could obtain the following descending order: LFP > ESA > MUA > SUA. It is important to note that the average inter-channel correlation of LFP and ESA above were obtained from recordings referenced to a single common reference (referred to as unipolar reference). To eliminate correlated noise commonly found in unipolar-based recordings, we applied a common average reference (CAR) referencing to the raw neural data. The CAR referencing significantly reduced the average inter-channel correlation for LFP (0.25 ± 0.10; 57.33%) but only marginally for ESA (0.45 ± 0.10; 0.2%).

**Fig. 3.**
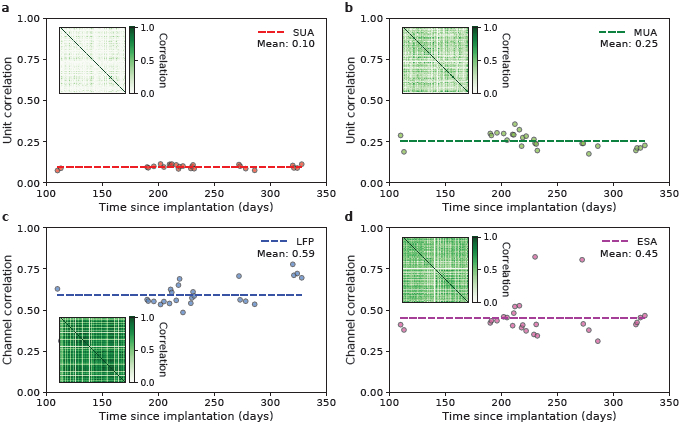
Inter-channel/unit correlation of neural signals over time. **a,b**, Average inter-unit correlation across all possible pairs of units within SUA and MUA, respectively. **c,d**, Average inter-channel correlation across all possible pairs of channels within LFP and ESA, respectively. The dashed lines represent the mean of correlation across all the sessions (*n* = 26). The insets depict the heatmap of inter-channel/unit correlation from different neural signals taken from the first session (I20160627_01).

### Decoding performance comparison across decoders

Next, we compared offline decoding of hand velocity from ESA signal using different decoders over time. The long-term decoding performance comparison results across ESA-driven decoders are depicted in Fig. 4. In the case of classical decoders, UKF yielded the highest average decoding performance (RMSE = 44.39 ± 1.38, CC = 0.82 ± 0.01; mean ± SEM), whereas KF yielded the lowest average decoding performance (RMSE = 58.69 ± 1.56, CC = 0.75 ± 0.01). In the case of deep learning decoders, QRNN produced the highest average decoding performance with RMSE of 35.48 ± 0.89 and CC of 0.89 ± 0.01, while SRNN produced the lowest average decoding performance with RMSE of 37.99 ± 0.99 and CC of 0.87 ± 0.01 (see Fig. 4a,b). Overall, the order of decoders from the highest to the lowest decoding performance was as follows: QRNN > LSTM > GRU > SRNN > UKF > WCF > WF > KF. As evident from Fig. 4c,e, all the deep learning decoders significantly outperformed all the classical decoders (two-tailed paired Wilcoxon signed-rank test; *** *p* < 0.001). By taking UKF, the best performing classical decoder, as the baseline, the deep learning decoders improved the average decoding performance by 13.91%–19.50% (RMSE) and 7.26%–8.90% (CC) as illustrated in Fig. 4d,f. In particular, our proposed QRNN decoder yielded decoding performance improvement on average by 19.50% ± 2.61% (RMSE) and 8.90% ± 2.27% (CC). Compared to the other classical decoders, QRNN achieved even higher decoding performance improvement. We found that QRNN yielded statistically significantly higher decoding performance than the other deep learning decoders (** *p* < 0.01, *** *p* < 0.001). We further sought to determine whether the superior decoding performance of the deep learning decoders over the classical decoders was consistently observed on each session. Our extensive results showed that all the deep learning decoders always outperformed their classical counterparts in each of 26 sessions for both RMSE and CC metrics as depicted in Fig. 4g and Fig. 4h, respectively. In addition, QRNN outperformed other deep learning decoders in 26/26 sessions (100% against SRNN), 25/26 sessions (96.15% against GRU), and 20/26 sessions (76.92% against LSTM).

**Fig. 4.**
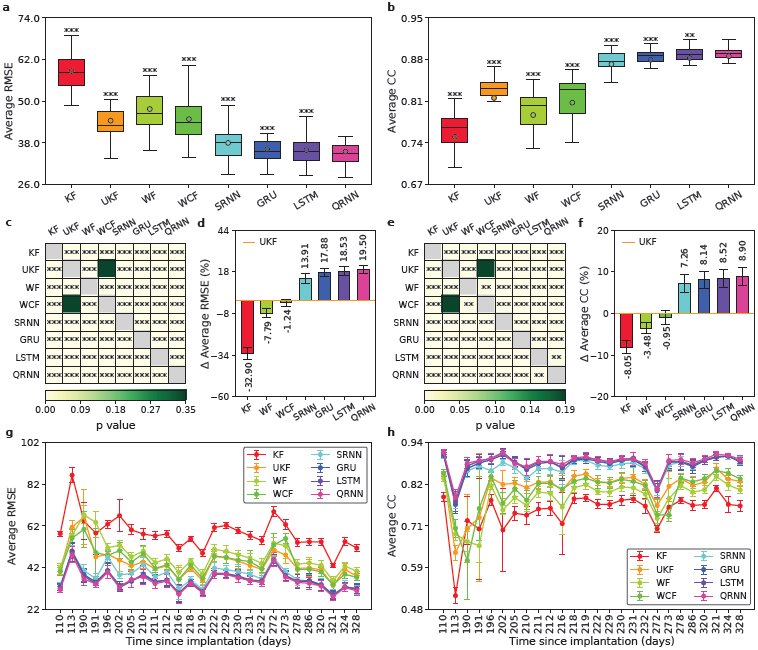
Long-term decoding performance comparison across different decoders driven by ESA signal. **a,b**, Boxplot comparison of long-term decoding performance across decoders as measured by average RMSE and CC, respectively. The horizontal lines inside the boxes denote the medians whereas the circles denote the means. Asterisks represent decoders whose decoding performance differed significantly from that of QRNN (two-tailed paired Wilcoxon signed-rank test; ** *p* < 0.01, *** *p* < 0.001). **c,e**, Heatmap of statistical significant matrix of decoding performance between all possible pairs of decoders as measured by average RMSE and CC, respectively. Each cell is coloured based on the *p*-value except cells in the main diagonal (grey; excluded from statistical test calculation). Asterisks indicate whether there were statistical significant differences between the pairs (two-tailed paired Wilcoxon signed-rank test; ** *p* < 0.01, *** *p* < 0.001). **d,f**, Average performance improvement/degradation (in %) of decoders with respect to that of UKF as measured by RMSE and CC metrics, respectively. Positive (negative) value indicates performance improvement (degradation). Black numeral texts represent the means whereas black error bars represent the 95% confidence intervals (*n* = 26). **g,h**, Decoding performance comparison over time across decoders in terms of average RMSE and CC, respectively. Coloured circles represent the means whereas coloured error bars represent the 95% confidence intervals (*n* = 10) for each session. The *x*-axes represent the number of days since electrode implantation. The *x*-axes’ ticks correspond to different sessions having irregular gaps between the consecutive sessions.

One may be interested in investigating whether above findings were also found in the cases of different neural signals. We therefore compared the decoding performance across decoders driven by SUA, MUA, and LFP. The decoding performance comparison results for the cases of SUA, MUA, and LFP are shown in Fig. S3, Fig. S4, and Fig. S5, respectively. We found consistent findings where all the deep learning decoders yielded significantly higher decoding performance than all the classical decoders (* *p* < 0.05, ** *p* < 0.01, *** *p* < 0.001). Moreover, we found that on average QRNN outperformed the other deep learning decoders across 3 neural signals, 26 sessions, and 2 metrics.

### Decoding performance comparison across neural signals

Further, we assessed the decoding performance of QRNN decoders across different neural signals. Fig. 5 presents the decoding performance comparison of QRNN decoders driven by LFP, SUA, MUA, and ESA. The results revealed that ESA yielded the highest average decoding performance with RMSE of 35.48 ± 0.89 and CC of 0.89 ± 0.01 (mean ± SEM). After ESA, the next highest average decoding performances (in descending order) were achieved by MUA (RMSE = 41.71 ± 0.86, CC = 0.84 ± 0.01), SUA (RMSE = 44.42 ± 1.01, CC = 0.81 ± 0.01), and LFP (RMSE = 47.45 ± 0.98, CC = 0.79 ± 0.01) as shown in Fig. 5a,b. Statistical tests showed that there were statistically significant differences between all possible pair combinations of neural signals (two-tailed paired Wilcoxon signed-rank test; *** *p* < 0.001) as illustrated in Fig. 5c,e. By taking SUA as the baseline, ESA improved the decoding performance by 20.12% ± 1.78 (RMSE) and 9.33% ± 1.52 (CC) as can be observed from Fig. 5d,f. The decoding performance improvements gained by ESA with respect to the other neural signals were as follows: 25.15% ± 2.63% (RMSE) and 12.99% ± 2.00% (CC) against LFP, and 15.02% ± 1.68% (RMSE) and 5.98% ± 0.80% (CC) against MUA. The superior decoding performance of ESA compared to the other neural signals was consistently observed across all 26 sessions (see Fig. 5g,h).

**Fig. 5.**
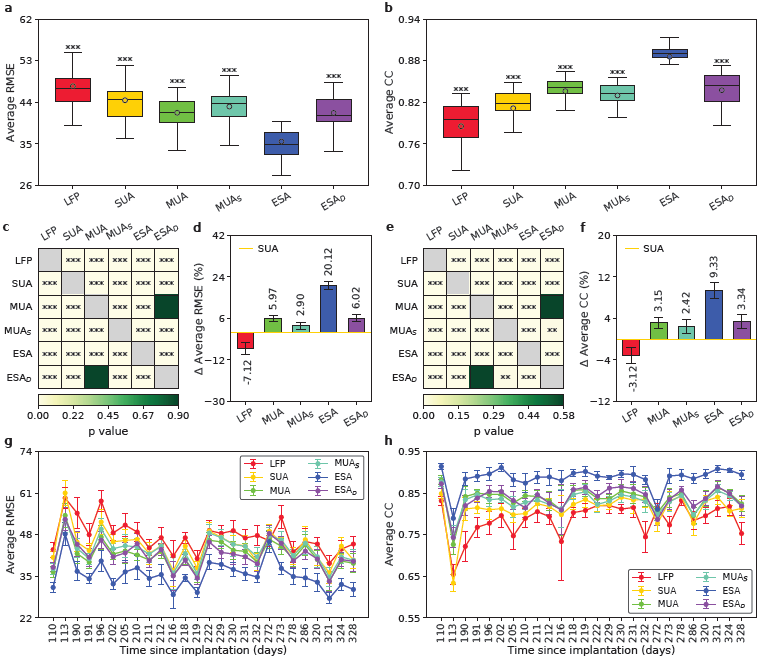
Long-term decoding performance comparison across QRNN decoders driven by different neural signals. **a,b**, Boxplot comparison of long-term decoding performance across different neural signals as measured by average RMSE and CC, respectively. The horizontal lines inside the boxes denote the medians whereas the circles denote the means. Asterisks represent neural signals whose decoding performance differed significantly from that of ESA (two-tailed paired Wilcoxon signed-rank test; *** *p* < 0.001). **c,e**, Heatmap of statistical significant matrix of decoding performance between all possible pairs of neural signals as measured by average RMSE and CC, respectively. Each cell is coloured based on the *p*-value except cells in the main diagonal (grey; excluded from statistical test calculation). Asterisks indicate whether there were statistical significant differences between the pairs (two-tailed paired Wilcoxon signed-rank test; ** *p* < 0.01, *** *p* < 0.001). **d,f**, Average performance improvement/degradation (in %) of neural signals with respect to that of SUA as measured by RMSE and CC metrics, respectively. Positive (negative) value indicates performance improvement (degradation). Black numeral texts represent the means whereas black error bars represent the 95% confidence intervals (*n* = 26). **g,h**, Decoding performance comparison over time across neural signals in terms of average RMSE and CC, respectively. Coloured circles represent the means whereas coloured error bars represent the 95% confidence intervals (*n* = 10) for each session. The *x*-axes represent the number of days since electrode implantation. The *x*-axes’ ticks correspond to different sessions having irregular gaps between the consecutive sessions.

To investigate whether the superior performance of ESA was dependence on the presence of spikes recorded on the same electrodes, we removed the spike components from ESA signal. We followed the procedure described in Stark and Abeles’ study,^53^ that is, by replacing all the detected spikes with high-frequency noise. This is done by subtracting smoothed spikes (computed through a five-sample moving average) from the associated raw neural signals. These spikeless ESA signals are referred to as despiked ESA (ESA_*D*_). We then performed the same offline decoding of hand kinematics using ESA_*D*_-driven QRNN decoder. The results revealed that although there was significant degradation (by 17.85% ± 2.20% in RMSE and 5.46% ± 0.62% in CC) in the average decoding performance relative to ESA, the average decoding performance of ESA_*D*_ was still significantly better than that of LFP and SUA (*** *p* < 0.001) and comparable to that of MUA, as shown in Fig. 5a,b and Fig. 5c,e). Specifically, the decoding performance improvements achieved by ESA_*D*_ were 12.02% ± 2.12% (RMSE) and 6.77% ± 1.57% (CC) against LFP, 6.02% ± 1.48% (RMSE) and 3.34% ± 1.32% (CC) against SUA, and 0.02% ± 1.38% (RMSE) and 0.18% ± 0.61% (CC) against MUA.

We also examined whether MUA inherently contained the same amount of information (i.e. spiking activity) as ESA and differed only from the signal processing point of view. We thus approximated the feature of ESA by convolving a Gaussian kernel with spike train from MUA. The smoothing parameter of the Gaussian kernel was selected by minimising RMSE between the actual ESA and approximated ESA (referred to as smoothed MUA or MUA_*S*_). We subsequently performed the same procedure of offline decoding using MUA_*S*_ -driven QRNN decoder. The results showed that the decoding performance of MUA_*S*_ was significantly lower (RMSE = 43.05 ± 0.85, CC = 0.83 ± 0.00) than that of MUA and ESA, indicating that MUA and ESA have a different amount of information content.

We extended the analysis of decoding performance comparison across neural signals using the other deep learning decoders. The comparison results for SRNN, GRU, and LSTM are presented in Fig. S6, Fig. S7, and Fig. S8, respectively. From these results, we found the same trend as above, that is, ESA produced statistically significantly higher decoding performance than LFP, SUA, and MUA (*** *p* < 0.001). The decoding performance of ESA corresponded to an improvement of 9.85%–30.24% (RMSE) and 4.76%–20.42% (CC) with respect to the other neural signals.

In order to examine the impact of different numbers of features (units or channels) on decoding performance, we selected *n* units or channels randomly from each neural signal. The *n* values used for SUA were {1, 10, 20, 30, …, 150}, while the *n* values used for LFP, MUA, ESA were {1, 10, 20, 30, …, 90}. We then compared the average decoding performance from *n* units/channels of each neural signal using QRNN decoder. We performed the same procedure for 30 iterations to obtain the mean and confidence interval. The comparison results from the first session (I20160627_01) across different neural signals are presented in Fig. 6a,b. We found that when *n* = 1, MUA yielded the highest decoding performance, followed by ESA, LFP, and SUA. However, when *n* > 1, ESA always produced the highest decoding performance compared to all the other neural signals. when *n* < 130, LFP showed better decoding performance than SUA but when *n ≥* 130, LFP was outperformed by SUA. As observed in Fig. 6a,b, the performance of all neural signals improved with the increasing *n* and reached a plateau after a certain value of *n*. ESA was found to reach a plateau quickest, whereas SUA reached a plateau slowest than the others. Similar findings were also observed when using data from the last session (I20170131_02) as displayed in Fig. S9. Snippet examples (10 s) of the true and decoded velocities (*x*-coordinate) from UKF and QRNN decoders driven by LFP, SUA, MUA, and ESA are graphically visualised in Fig. 6c-f. Likewise, snippet examples of the true and decoded velocities in *y*-coordinate from the same decoders are plotted in Fig. 6g-j.

**Fig. 6.**
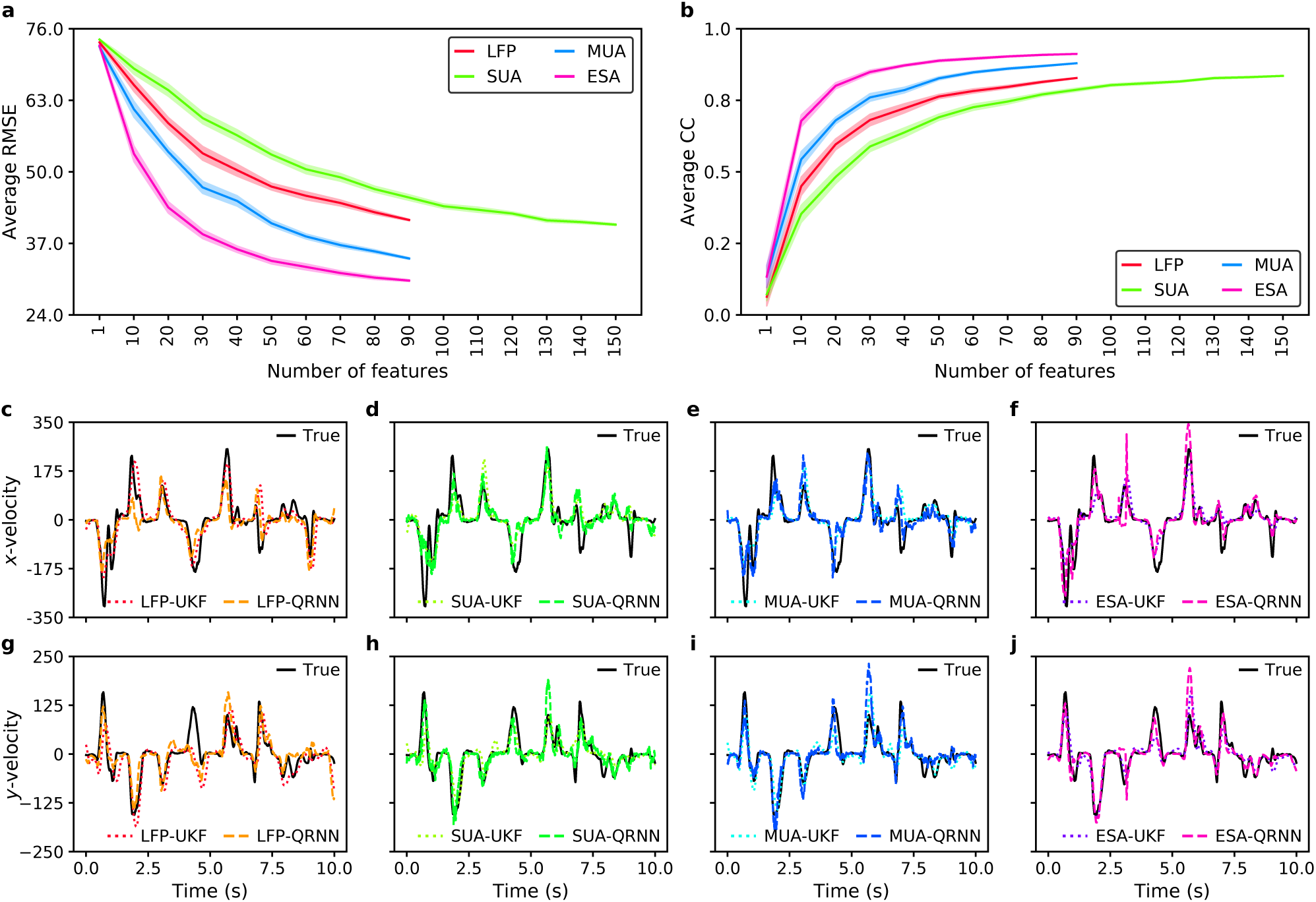
Decoding performance comparison across different types of neural signals. **a,b**, Impact of different number of features (units or channels) on the decoding comparison of QRNN decoders driven by different neural signals as measured by average RMSE and CC, respectively. The shaded areas represent the 95% confidence intervals (*n* = 30). **c-f**, Snippet examples of true and decoded *x*-velocities from LFP-, SUA-, MUA-, and ESA-driven decoders, respectively. **g-j**, Snippet examples of true and decoded *y*-velocities from LFP-, SUA-, MUA-, and ESA-driven decoders, respectively. Data are taken from the first session (I20160627_01).

### Impact of different training schemes and the amount of training data

To investigate how frequently a decoder should be updated or whether a static decoder could sustain high decoding performance over time, we trained QRNN decoder using three training schemes: (1) trained the decoder only at the first session and kept it fixed for the rest sessions (referred to as fixed QRNN), (2) retrained the decoder at every session without taking into account the data from the previous sessions (referred to as retrained QRNN), and (3) retrained the decoder at every session by resuming (i.e. continuing) the training from the previous session (referred to as resumed QRNN). The training was performed on 80% of data from each session, while the testing was performed on 10% of data sequentially located after the training data to mimic the real application scenario. The results showed that, on average, the retrained QRNN yielded slightly higher decoding performance than the resumed QRNN and significantly higher decoding performance than the fixed QRNN as illustrated in Fig. 7a,b. Although the fixed QRNN exhibited degraded performance compared to the retrained QRNN and resumed QRNN, its decoding performance was still moderately good except at 3 out of 26 sessions.

**Fig. 7.**
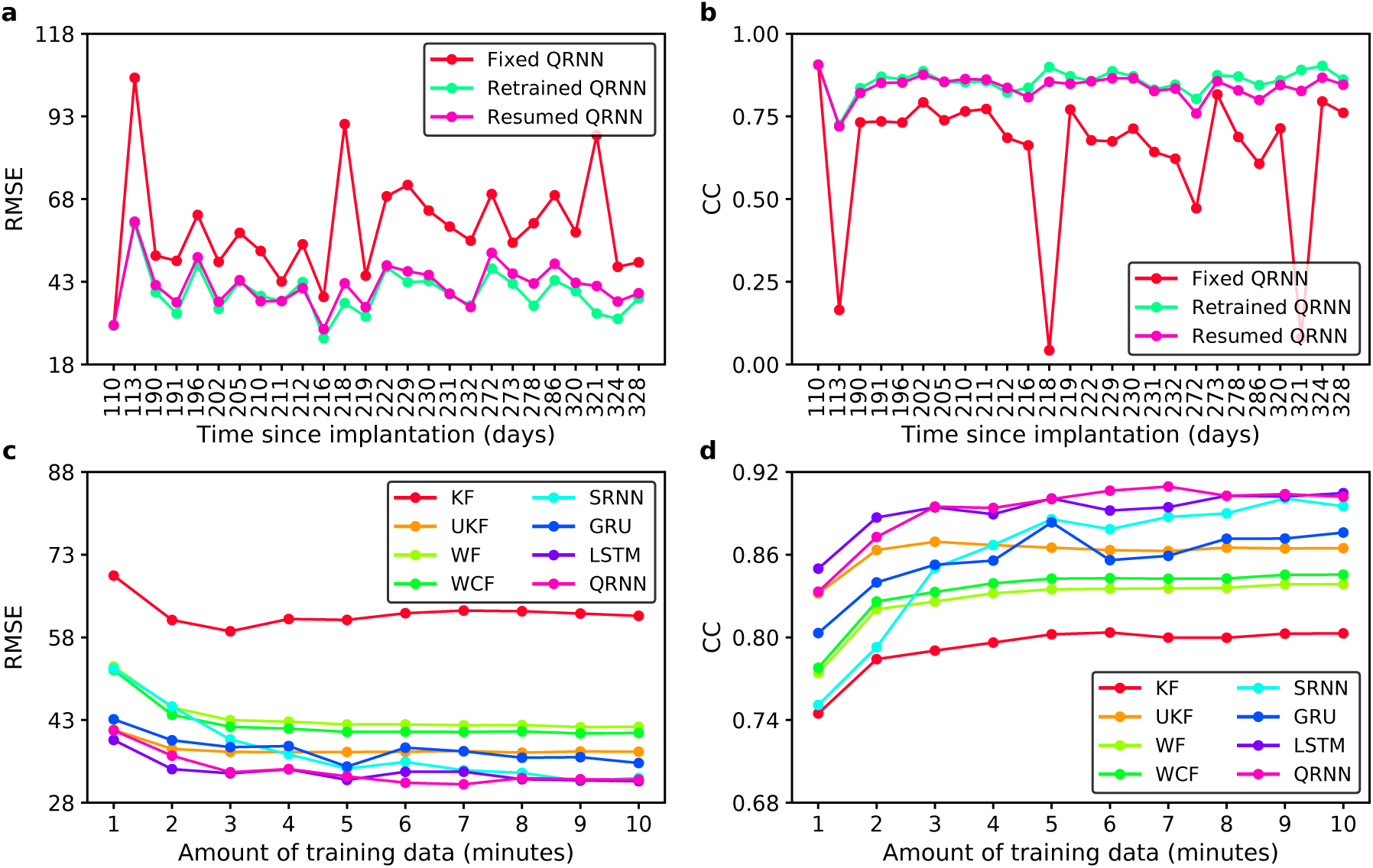
Decoding performance comparison across decoders under different training schemes and amount of training data. **a,b**, Impact of the training schemes on long-term decoding performance of QRNN decoder across sessions as measured by RMSE and CC, respectively. **c,d**, Impact of the amount of training data on decoding performance of different decoders as measured by RMSE and CC, respectively. Data are taken from the first session (I20160627_01).

The high accuracy of deep learning is often attributed to a large amount of data it uses during training. To test this preconception, we varied the amount of training data from 1 to 10 minutes with an increment of 1 minute. We analysed and compared the decoding performance of deep learning decoders with that of the classical decoders. Fig. 7c and Fig. 7d plot the impact of increasing the amount of training data on the decoding performance across eight different decoders measured by RMSE and CC, respectively. We found that, on average, the decoding performance of all decoders improved the increasing amount of training data. However, all the decoders on average had already reached a plateau when the training data equals to 5 minutes. Another finding was that both QRNN and LSTM outperformed UKF (the best performing classical decoder) even when the training data was small (*≤* 3-minute length). On the other hand, SRNN and GRU were outperformed by UKF when using a small amount of training data. Regardless of the amount of training data, KF was always found to yield the lowest decoding performance.

## Discussion

Previous studies have shown that SUA-based BMIs yielded good decoding performance in acute settings.^35, 36, 54^ The fact that SUA signals fade over time makes it less desirable chronically and poses a major challenge towards robust and accurate BMIs. To address this issue, here, we introduce a signal/decoder co-design methodology by exploiting the synergism between the input signal and decoding algorithm within the design development process. Specifically, we propose simultaneously an alternative input signal based on entire spiking activity (ESA) and a new decoding algorithm based on quasi-recurrent neural network (QRNN). We comprehensively evaluated its decoding accuracy and robustness over a long period of time. Firstly, we characterised the long-term stability and quality of ESA and compared its characteristics with commonly used neural signals which include SUA, MUA, and LFP. As expected, SUA showed statistically significant decrease in all three characteristics over time: (1) number of recorded units, (2) average peak-to-peak amplitude (Vpp; in *µ*m), and (3) mean spike rate (Hz). MUA was found to exhibit a significant decrease only in the mean spike rate, indicating its better stability than SUA. On the other hand, LFP did not show a decreasing trend in the mean amplitude, while ESA showed a slow (not statistically significant) decreasing trend, suggesting that both LFP and ESA are more stable over time than SUA and MUA. One approach to mitigate the signal instability issue faced by SUA-based BMIs is by utilising MUA as an alternative input neural signal.^25, 29, 30, 36^ Another approach is to use LFP, which has been believed and demonstrated to be stable over a long period of time even when the spikes have already deteriorated or lost.^19, 24, 39, 55, 56^ Our results confirm the previous studies by showing that both LFP and MUA exhibit better signal stability than SUA. We further extend the previous findings by demonstrating that ESA also possesses the characteristic of long-term signal stability, which could potentially be used as an alternative input neural signal for BMIs.

To examine the redundancy characteristics of neural signals, we compared the inter-channel/unit correlation between all possible pair combinations of channels/units within each neural signal. LFP was found to have the highest average correlation with a significantly large gap from the other neural signals, while SUA was found to have the lowest average correlation. This is in good agreement with prior studies.^34, 36, 39, 57^ Overall comparison results on long-term data yielded the following order (sorted descendingly): LFP > ESA > MUA > SUA. The results may also indicate the degree of independence between channels/units within each neural signal. The high redundancy of LFP might be attributed to the contamination of correlated noise arising from different sources, e.g. non-silent reference and volume conduction^58, 59^ as commonly observed in LFP recordings with unipolar reference. To remove the correlated noise, we re-referenced the raw neural signals using common average reference (CAR) and re-computed the LFP. The resulting CAR-based LFP had substantially lower inter-channel correlation. In addition, the different level of redundancy could be related to the spatial integration or coverage of each neural signal. LFP is thought to mainly reflect summed synaptic activity from a local neuronal population ranging from a radius of a few hundred micrometres to a few millimetres around the recording electrodes.^60–63^ As the Utah array has an inter-electrode distance of 400 *µ*m, the recorded LFP on each electrode could therefore contain (to some degree) overlapping neural activity. On the other hand, SUA represents spike (action potential) activity from individual neurons which can be well discriminated within a radius less than ∼50 *µ*m^64^ or up to ∼130 *µ*m^65^ from the recording electrodes. The small spatial coverage of SUA may partly explain its low inter-unit correlation and high degree of independence. MUA, which represents the aggregate spikes from an ensemble of neurons near the electrodes, has a spatial coverage larger than SUA but less than LFP. Several studies reported that the spatial coverage of MUA is within a radius of ∼140–300 *µ*m.^66–69^ As for ESA, it is considered to reflect an instantaneous measure of the number and size of spikes from a population of neurons around the electrodes.^70, 71^ The spatial coverage of ESA is likely larger than that of MUA (but smaller than that of LFP) as it includes the contribution from small amplitude spikes (either from small nearby neurons or big farther neurons) which may not be detected through threshold crossing technique.^72, 73^

Next, using ESA as the input, we benchmarked offline decoding performance of our proposed QRNN decoder against three deep learning decoders (SRNN, GRU, and LSTM) and four classical decoders (WF, WCF, KF, and UKF). Our analysis resulted in several findings as follows. First, QRNN yielded significantly higher decoding accuracy compared to all the other decoders and could maintain its high accuracy over a long period of time. Second, all the deep learning decoders significantly outperformed all the classical decoders. Third, the superiority of the deep learning decoders, particularly QRNN, over the classical counterparts were consistently observed across different neural signals (SUA, MUA, and LFP) and sessions. Forth, within the category of classical decoders, UKF emerged as the highest performing decoder. These findings reinforce previous studies highlighting the high performance of deep learning decoders (SRNN, GRU, LSTM) in comparison to classical decoders.^49–52^ This study further expands the previous studies by demonstrating that QRNN yielded better decoding accuracy and robustness over time than the other deep learning decoders. There is concern that the high performance of deep learning decoders is attributable to a large enough amount of training data. To test this preconception, we varied the amount of training data from 1 to 10 minutes and compared the decoding performance across deep learning and classical decoders. Comparison results demonstrated that all the decoders exhibited performance decline with the decreasing amount of training data. However, we found that QRNN and LSTM still performed considerably good compared to the others even when the training data were small. This finding is in line with a recent report by Glaser *et al*.^50^ One potential implication of this finding is that the data collection for decoder training could be shortened which in turn speeds up the decoder calibration and deployment processes while minimising the subject’s learning fatigue. Due to the variability and nonstationarity of neural signals, a fixed (i.e. static) decoder may degrade over the course of hours, days, or months. To address this issue, the decoder needs to be recalibrated regularly at the beginning of each session or when there is a significant drop in the decoding performance. Here, we investigate the impact of three training schemes on the decoding performance: retraining only at the first session (fixed decoder); retraining from scratch at every session (retrained decoder); and resuming the training from the previous session (resumed decoder). Extensive results using different deep learning decoders showed that retrained decoders performed slightly better than resumed decoders and substantially better than fixed decoders. Taking into account the information from previous sessions during training as in resumed decoders on average did not improve the decoding performance compared to retraining from scratch at every session (retrained decoders). This could indicate the disparity between the previous and current sessions arising probably from changes in neural encoding pattern or recording degradation.^24, 74^ Even though fixed decoders exhibited a significant performance decline (as anticipated), their decoding performance was still moderately good in most (except a few) sessions. This suggests that under circumstances where decoder recalibration cannot be carried out frequently on per-session basis, fixed decoder can be leveraged. The recalibration process can then be performed only when the performance of a fixed decoder substantially deteriorates.

Further, we quantified movement-related information content within each neural signal by comparing their decoding performance using QRNN decoder. Results revealed that ESA significantly outperformed the other neural signals and could sustain its high decoding performance across all the sessions. The order of neural signals according to their decoding performance (from high to low) was ESA > MUA > SUA > LFP. The same trend was consistently observed when using different deep learning decoders, indicating that ESA holds rich information about the movement parameters. To gain more insights into whether the superiority of ESA is dependence on the presence of spikes recorded from the same electrodes, we removed the spike components from the raw neural signals and recomputed ESA (termed as depsiked ESA or ESA_*D*_) using a method described by Stark and Abeles.^53^ We found that ESA_*D*_ still produced better decoding performance than SUA, MUA, and LFP, suggesting that ESA contains additional information other than spikes detected through threshold crossing technique (as in SUA and MUA). Spike detection through threshold crossing technique relies on manual or automatic selection of threshold value. This could be problematic in chronic recordings with low SNR or high variability over time. If the threshold value is too small, it may detect some background noise as spikes, increasing the false positive rate. Conversely, too high threshold value could leave true spikes undetected. In addition, threshold crossing technique may provide a biased estimate of spikes in favour of large neurons (with large amplitude), thus leaving spikes from small neurons undetected.^72, 73^ Contrary to SUA and MUA, ESA uses a simple, automated, and threshold-less method that allows a more reliable and less biased estimate of spikes and takes into account the contribution from all neurons (including the small ones) in the vicinity of the electrodes.^72, 73^ We examined this issue further by approximating ESA from MUA (referred to as smoothed MUA or MUA_*S*_) through a Gaussian kernel-based smoothing with an optimised width parameter. Results showed that MUA_*S*_ could not approach the level of decoding performance of ESA. This may support the idea that there is additional information in ESA which is not available in MUA.

Another finding in our study is that LFP yielded the lowest decoding performance despite its better long-term signal stability, indicating its less movement-related information content than the other neural signals. This finding is consistent with a considerable number of prior studies.^21, 33, 38, 39, 75^ It is worth noting that this observation was obtained from the same cortical depth due to the fixed geometry of Utah array (1 mm electrode length), which is perhaps not the ideal location for LFP recordings. It has been shown that the properties of LFP recordings depend on the recording depth.^76^ Therefore, this finding may not be generalisable to data recorded using linear (variable-depth) electrodes. In the long run, as the number of detected spikes decreases, LFP could outperform both SUA and MUA as demonstrated in previous reports by Flint *et al.*^19, 44^ and Wang *et al*.^24^

In addition, our analysis on unit/channel-dropping curve showed that LFP yielded higher decoding performance than SUA when the number of channels/units were relatively small. When using all the available channels/units, SUA outperformed LFP, which compares well with previous works.^33, 34, 38, 39^ This could act as an additional indication that LFP is more redundant (less independence) than SUA. On the other hand, ESA always yielded the highest decoding performance compared to the other neural signals across different decoders, sessions, and numbers of channels. With the increasing number of channels/units, the performance of ESA reached a plateau quicker than the others, suggesting that ESA exhibits some degree of redundancy. The implication of this is that we could obtain accurate and robust decoding performance using ESA even without using all the available channels, which can reduce computational complexity, memory footprint, and power consumption.

ESA has been previously used to measure evoked responses in the visual cortex upon receiving various visual stimuli,^72, 73, 77, 78^ whereas QRNN has been mainly used for natural language processing (NLP) applications.^79, 80^ There is no previous study that uses both ESA and QRNN for intracortical BMI application. To the best of our knowledge, our study is the first to systematically and comprehensively compare long-term signal and decoding stability across different neural signals and decoders simultaneously. Our results agree well with prior studies that used a neural signal which in principle is similar to ESA.^53, 81^ However, there are differences in the signal processing point of view to obtain such signal. After high-pass filtering, this study uses absolute operation (i.e. full-wave rectification) followed by a low-pass filtering at 12 Hz, whereas those prior studies use root mean square (RMS) operation followed by a low-pass filtering at 100 Hz. Absolute operation is simpler and easier than RMS operation especially when implemented on hardware (i.e. chip). Lower frequency cut-off enables signals to be downsampled into a lower sampling rate, which in turn reduces the computational complexity.

The limitation of this study is that we use neural data recorded approximately from 3.6 to 10.9 months after implantation, corresponding to a total of 26 sessions. Although there is a gradual decline over time, the number of detected spikes at the last session is still relatively high. It remains to be tested whether ESA-based QRNN decoder could maintain its high decoding performance for many years when spikes have severely diminished or even lost. Our analysis using despiked ESA suggests that ESA could still provide good performance even when the spikes are lost. A recent study using a signal similar to ESA demonstrated that such a signal could sustain good performance over three years worth of recordings.^81^

Taken together, our overall results demonstrate that ESA signal coupled with deep learning decoder, particularly QRNN, yields chronically robust and exceptionally accurate decoding performance over time. Furthermore, the simple, automated, threshold-less processing requirements combined with the low bandwidth of ESA address key challenges in the implementation of a fully implantable wireless BMI system.

## Methods

### Neural recordings

Neural recordings were obtained from a public dataset deposited by Sabes lab.^82^ The neural data were recorded from primary motor cortex (M1) area of an adult male Rhesus macaque monkey (*Macaca mulatta*), indicated by I. The recordings were made by using a 96-channel Utah microelectrode array (platinum contact, 400 kΩ nominal impedance, 400 *µ*m interelectrode spacing, 1 mm electrode length) referenced to a silver wire placed under the dura (several cm away from the electrodes). The recordings were preamplified and filtered by using a 4th-order lowpass filter at 7.5 kHz with a roll-off of 24 dB per octave. The recordings were then digitised with 16-bit resolution at 24.4 kHz sampling rate. These digitised recordings are hereinafter referred to as raw neural signals. Details of the experimental setup including the subject, surgical procedure, and recording setup have been described elsewhere.^83^ The neural data used in this work consist of 26 recording sessions spanning 7.3 months between the first (I20160627_01) and last (I20170131_02) sessions. The first and last sessions correspond to 110 and 328 days since the electrode implantation, respectively. The recording duration varies from 6 to 13.6 minutes with an average of 8.88 ± 1.96 minutes (mean ± SD).

### Behavioural task

The behavioural task has been previously described in detail elsewhere.^83^ Briefly, the monkey was trained to perform a point-to-point task, that is, to reach randomly drawn circular targets which were uniformly distributed around an 8 *×* 8 square grid, as illustrated in Fig. 1. The target was acquired when the monkey reached the target using his fingertip and held it for 450 ms. Upon every target acquisition, a new random target was presented immediately without an inter-trial interval. The fingertip position of the reaching hand and the target position (in x,y Cartesian coordinates with mm unit) were both sampled at 250 Hz. The position data were then low pass filtered with a non-causal, 4th-order Butterworth filter at 10 Hz to reject sensor noise. Velocity and acceleration data were computed by using first and second derivative of the position data.

### Signal processing and feature extraction

From the raw neural signals, we computed four different types of neural signals, namely local field potential (LFP), single-unit activity (SUA), multiunit activity (MUA), and entire spiking activity (ESA). Examples of these four neural signals along with their associated features are shown in Fig. 1. The signal processing and feature extraction algorithms were all implemented in Python and are described briefly as follows.

#### Local field potential (LFP)

The raw neural signals were first digitally re-referenced using common average reference (CAR) to eliminate common noise. LFPs were obtained by low-pass filtering the re-referenced neural signals with a 4th-order Butterworth filter at 300 Hz followed by downsampling them to 1 kHz. To avoid any phase shift, the filtering processes in this work were by default performed on the forward and backward directions (unless otherwise stated). Local motor potential (LMP), a time-domain average amplitude feature, was extracted by using a moving average filter with 256 ms rectangular window. The LMP feature was extracted through an overlapping fashion (252 ms overlap) to yield a sample every 4 ms, thereby matching the timescale of the kinematic data. The LMP was chosen because it has been shown to yield better decoding performance than other LFP features.^21, 84^ LFP features from all the 96 channels were included in our experiments and analyses.

#### Single-unit activity (SUA)

The raw neural signals were band-pass filtered with a causal, 4th-order Butterworth IIR filter between 500 to 5000 Hz. Spikes were then detected whenever the absolute value of the band-passed signals crossed a threshold value (typically set to between 3.5 and 4.0 times the standard deviation). SUA was obtained by sorting (i.e. classifying) the detected spikes into distinct putative single units via principal component analysis and template matching. This means each channel could contain more than one SUA. More detailed information on the spike detection and sorting processes can be found in.^83^ We computed spike rate from SUA using a 256 ms rectangular window sliding every 4 ms. We Only included SUAs with spike rates above 0.5 Hz, which resulted in a varying number of SUA ranging from 91 to 157 units with an average of 125.73 ± 13.95 units (mean ± SD) across 26 recording sessions.

#### Multi-unit activity (MUA)

In this study, MUA is defined as all the spikes detected through threshold crossing technique. To obtain MUA, we performed spike detection (as described in the above SUA processing) but without spike sorting. Thus, for each channel, we obtained a maximum of one MUA. We then computed spike rate from MUA using a 256 ms rectangular window sliding every 4 ms. Similar to that of SUA, only MUAs with spike rates exceeding 0.5 Hz were included. This yielded a number of MUA that varied from 77 to 95 units with an average of 87.08 ± 4.44 (mean ± SD) across 26 recording sessions.

#### Entire spiking activity (ESA)

The term ESA was firstly used by Drebitz *et al.* in 2018^85^ but its underlying principle has been known and used for several decades.^70, 86^ ESA, sometimes referred to as the “envelope of the spiking activity”, was previously utilised to measure evoked responses (e.g. receptive field) in the visual cortex of monkeys upon receiving various types of visual stimuli.^72, 73, 77, 78, 87^ While both SUA and MUA are represented by a sequence of binary signals, ESA is represented by a continuous signal reflecting an instantaneous measure of the number and size of spikes from a population of neurons surrounding the electrode tip.^70, 71^ ESA was obtained by digitally re-referencing the raw neural signals with common average reference (CAR) and followed by high-pass filtering with 1st-order Butterworth filter at 300 Hz. The filtered signals were then full-wave rectified (i.e. taking the absolute value), low-pass filtered with 1st-order Butterworth filter at 12 Hz and downsampled to 1 kHz. We extracted a time-domain average amplitude feature (similar to LMP) using a moving average filter with a 256 ms rectangular window overlapping by 252 ms (yielding a feature sample every 4 ms).

### Decoding methods

We benchmarked eight decoding algorithms which can be categorised into classical decoders and deep learning decoders; each category consists of four decoding algorithms. The classical decoders include Wiener filter (WF), Wiener cascade filter (WCF), Kalman filter (KF), and unscented Kalman filter (UKF), while the deep learning decoders include standard recurrent neural network (SRNN), long short-term memory (LSTM), gated recurrent unit (GRU), and quasi-recurrent neural network (QRNN). TensorFlow (Keras) deep learning framework was used to implement the deep learning decoders. Each of these eight decoders is briefly described as follows.

#### Wiener filter (WF)

WF, introduced by Norbert Wiener in 1940s, is a linear filter used to estimate an unknown signal of interest (e.g. kinematic data) from noisy observations (e.g. neural data) under the assumption of known stationary signal and additive noise. WF produces an optimal estimate of the desired signal in the sense of minimum mean squared error. WFs were used as a decoding algorithm in several prior BMI studies.^5, 40, 41^ In this study, the number of taps (i.e. lags) of WF was empirically set to 15. The weights of the WF were estimated by using linear least squares with *ℓ*_2_ (also known as ridge or Tikhonov) regularisation. The regularisation parameter (*α*) was set to 0 (by default) and increased to a large value if there was a significant drop in decoding performance.

#### Wiener cascade filter (WCF)

WCF is an extension of WF to deal with nonlinear dynamic systems. It is composed of two stages: a linear dynamic system followed by a static nonlinearity.^19, 39, 44^ The first stage was implemented by a WF, whereas the second stage was implemented by fitting a 3rd order polynomial between the WF output and the kinematic data. The number of taps and the regularisation parameter (*α*) were set equal to that of the WF.

#### Kalman filter (KF)

KF, primarily developed by Rudolf E. Kalman in 1960s, extends WF to nonstationary signals. It is one of the most popular estimation algorithms and has been employed as a decoding algorithm in numerous BMI studies.^6, 9, 32, 33, 42^ KF is an optimal estimator for linear dynamic systems with linear measurement models assuming all noise is Gaussian. KF algorithm consists of two steps, *prediction* and *update*, which are performed in a recursive manner. Firstly, KF predicts the current states and their uncertainties from the previous states. These predicted states are then updated by combining them with the new measurement using a weighted average. We followed the KF implementation described in Wu *et al.*’s study^42^ and added *ℓ*_2_ regularisation to the estimation of state transition and measurement matrices. The regularisation parameter (*α*) was by default set to 0 except in a few sessions (increased to a large value) when a significant drop in decoding performance was encountered.

#### Unscented Kalman filter (UKF)

UKF is an extension of KF to deal with nonlinear systems. It utilises an unscented transform, that is, a method to calculate the statistics of a random variable passing through a nonlinear transformation. It builds on the intuition that approximating a probability distribution is easier than approximating an arbitrary nonlinear function.^88^ This can be accomplished by using a minimal set of carefully and deterministically chosen sample points known as sigma points. We followed the UKF implementation described in Li *et al.*’s study.^45^ The state transition and measurement matrices were estimated using the same method as of the KF. The scaling parameter (*κ*) and number of taps were both set to 1.

#### Standard recurrent neural network (SRNN)

SRNN is a class of neural networks where the outputs from the previous timestep are used as inputs in the current timestep. SRNN has a “memory” in the form of a “hidden state” which allows it to remember sequential information that has been calculated so far. SRNN offers several advantages, including ability to process variable length input, taking into account historical information, and shared weights across timesteps. The output at timestep *t* in SRNN can be formulated as:

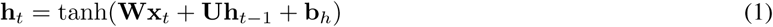

where **x**, **h**, and tanh denote the input, hidden state, and hyperbolic tangent activation function, respectively. **W**, **U** and **b** represent input weight, recurrent weight, and bias vector. The graphical illustration of SRNN is shown in Fig. S1a.

#### Long short-term memory (LSTM)

LSTM, proposed by Hochreiter and Schmidhuber in 1997,^89^ addresses the vanishing/exploding gradient problem encountered when training SRNNs. LSTMs have become one of the most popular deep learning architectures and have achieved state-of-the-art performance in a wide range of machine learning problems, especially those dealing with time-series data.^90^ LSTMs can effectively learn long-term temporal dependencies via a memory cell that maintains its state overtime and a gating mechanism that controls the flow of information into and out of the memory cell. We employed a commonly used variant of LSTM architectures which is formulated as follows:

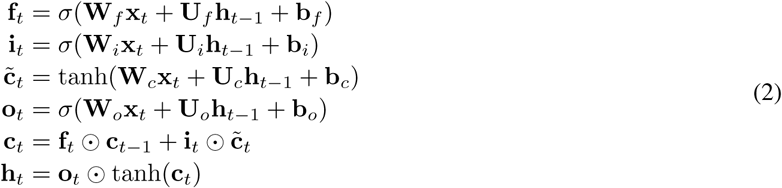

where **x**, **h**, **f**, **i**, **o**, **c** consecutively represent the input, output, forget gate, input gate, output gate, and memory cell. *σ* and ⊙ denote the logistic sigmoid activation function and element-wise multiplication operator. The graphical illustration of LSTM is shown in Fig. S1b.

#### Gated recurrent unit (GRU)

GRU is another variant of recurrent neural networks (RNNs) proposed by Cho *et al.* in 2014^91^ that uses a gating mechanism similar to LSTM. It has less parameters and less structural complexity than LSTM since it combines the forget gate and input gate into a single update gate and combines the memory cell and hidden state. Each component of GRU is defined by the following equations:

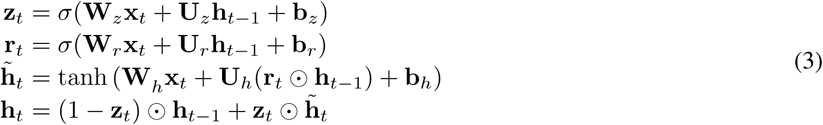

where **z**, **r**, **h** represent the update gate, reset gate, and hidden state, respectively. Chung *et al.* demonstrated empirically that the performance of GRU was comparable to that of LSTM across three sequence modelling tasks.^92^ GRU is graphically illustrated in Fig. S1c.

#### Quasi-recurrent neural network (QRNN)

QRNN was recently introduced by Bradbury *et al.* in 2016^79^ that combines the parallelism power of convolutional neural networks (CNNs) with the capability of RNNs in learning temporal dependencies of sequential data. It is composed of two components: convolutional component that performs convolutions in parallel across timesteps and pooling component that handles temporal dependencies in parallel across feature dimensions. The full set of equations in QRNN is defined as follows:

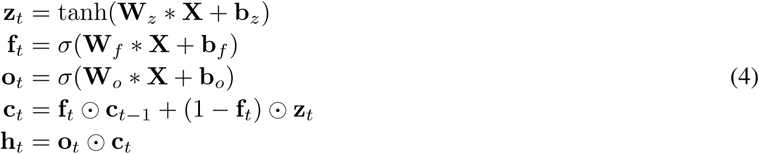

where **z**, **f**, **o**, **c**, **h** consecutively represent the candidate vectors, forget gate, output gate, memory cell, and hidden state. Operator ∗ denotes a masked convolution which is a type of convolutions that depends only on the past and present inputs (i.e. cannot access inputs from future timesteps). QRNN is graphically illustrated in Fig. S1d.

### Hyperparameter optimisation

For each deep learning decoder, the final output was obtained by connecting the last timestep of output from the RNN layer with a fully connected layer. The number of layers and timesteps were empirically set to 1 and 2, respectively. Other hyperparameters such as the number of units, number of epochs, batch size, dropout rate and learning rate were determined through hyperparameter optimisation.

We split each recording session into 10 non-overlapping contiguous segments of equal size which were then categorised into three different sets: training set (8 concatenated segments), validation set (1 segment) and testing set (1 segment). The training set was used to train the deep learning (DL) decoders; the validation set was used to optimise the hyperparameters of the DL decoders; the testing set was used to evaluate the performance of the optimised DL decoders. All features being fed to the DL decoders were standardised (i.e. z-transformed) to have zero mean and unit variance. The DL decoders were trained using a root mean squared error (RMSE) loss function and an RMSprop optimiser. Bayesian optimisation library called Hyperopt^93^ was used for the hyperparameter optimisation which was conducted separately for each type of neural signal and run for 300 iterations. To save the computational time of experiments, the hyperparameter optimisation was performed only once using the first recording session. The hyperparameters for each DL decoder were optimised from predefined hyperparameter search spaces listed in Table S1. The value ranges for number of units were specified so that the minimum and maximum possible number of parameters (weights) are comparable across different decoders. The hyperparameter optimisation result for the case of ESA-driven QRNN decoder is visually illustrated in Fig. S2. The resulting optimised hyperparameters for deep learning decoders driven by ESA and for QRNN decoder driven by LFP, SUA, and MUA are provided in Table 1. The optimised hyperparameters for deep learning decoders driven by LFP, SUA, and MUA are given in Table S2.

**Table 1.**
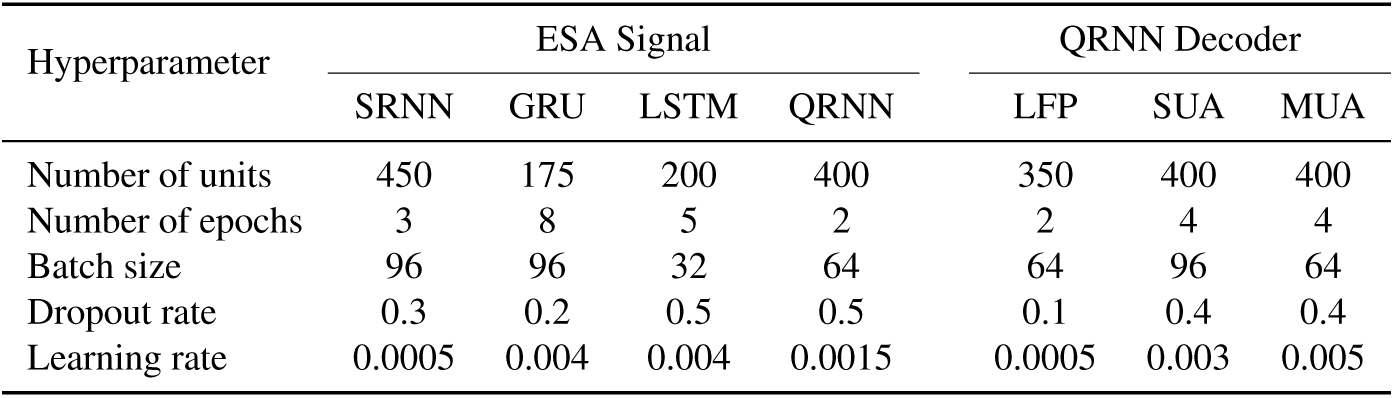
Hyperparameter configurations of deep learning decoders driven by ESA and QRNN decoders driven by other neural signals.

### Decoder training schemes

According to the training scheme being used, we divided the decoders into three categories: (1) fixed decoder, (2) retrained decoder, and (3) resumed decoder. Each of these categories is briefly described as follows.

#### Fixed decoder

In this category, the training was performed at the first session and the trained weights were kept fixed for the rest of the sessions. If in the subsequent sessions there were channels that did not record neural activity, these channels were assigned with zeros to keep the input structure (i.e. the number of channels) remain the same.

#### Retrained decoder

In this category, the training was performed at every session without taking into account the information from the previous session. All the decoders were by default trained by using this scheme (unless otherwise stated).

#### Resumed decoder

In this category, the training was performed at every session while keeping the information from the previous session. This is done by storing the model (i.e. decoder) from the previous session and resuming the training from where we left off using the current session data.

### Performance evaluation and metrics

The decoding performance was evaluated using two commonly used metrics: (1) root mean square error (RMSE) and (2) Pearson’s correlation coefficient (CC).^32, 33, 38^ RMSE represents the average magnitude of the decoding error; CC which is a translation and scale invariant metric assesses the linear correlation (shape similarity) between the true and decoded hand kinematics. RMSE and CC are formulated as follows:

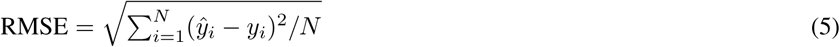

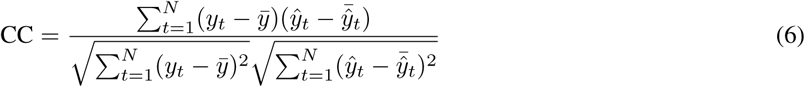

where *y*_*t*_ and *ŷ*_*t*_ denote the true and decoded hand kinematics at timestep *t*, respectively, and *N* represents the total number of samples.

### Statistical analysis

The stability of each neural signal over time was assessed by calculating the slope of its regression line. The slope was calculated using Huber regression, one type of robust linear regression methods, that is less sensitive to outliers without ignoring their effect. We tested whether the neural signal was statistically significantly stable using a one-tailed one-sample *t*-test. For each session, the decoding performance was evaluated on 10 different segments within the testing set to obtain the mean and confidence interval. To test whether there was statistical difference between a pair combination of neural signal and decoder, we used a Wilcoxon signed-rank test. The significance level (*α*) was set to 0.05.

Boxplots were used to visualise the decoding performance comparison among neural signals/decoders across 26 sessions. The horizontal line and circle mark inside each box represent the median and mean, respectively. The coloured solid box represents the interquartil range (between 25th and 75th percentiles). The whisker extends 1.5 times the interquartile range. All the analyses were conducted in Python.

## Data availability

Data are available from Zenodo at https://zenodo.org/record/583331.

## Acknowledgments

This work was supported by the UK Engineering and Physical Sciences Research Council (grant number EP/M020975/1) and Indonesia Endowment Fund for Education (LPDP) graduate scholarship program (grant number PRJ-123/LPDP/2016). The authors thank J. E. O’Doherty, M. M. B. Cardoso, J. G. Makin and P. N. Sabes for making their data publicly available.

## Author contributions

N.A., T.G.C. and C.S.B. conceived the study, N.A. developed the methodology, conducted the experiments, analysed and visualised the data, N.A., T.G.C and C.S.B interpreted the results, N.A. wrote the original draft. All authors reviewed and edited the manuscript.

## Supplementary Information

**Fig. S1.**
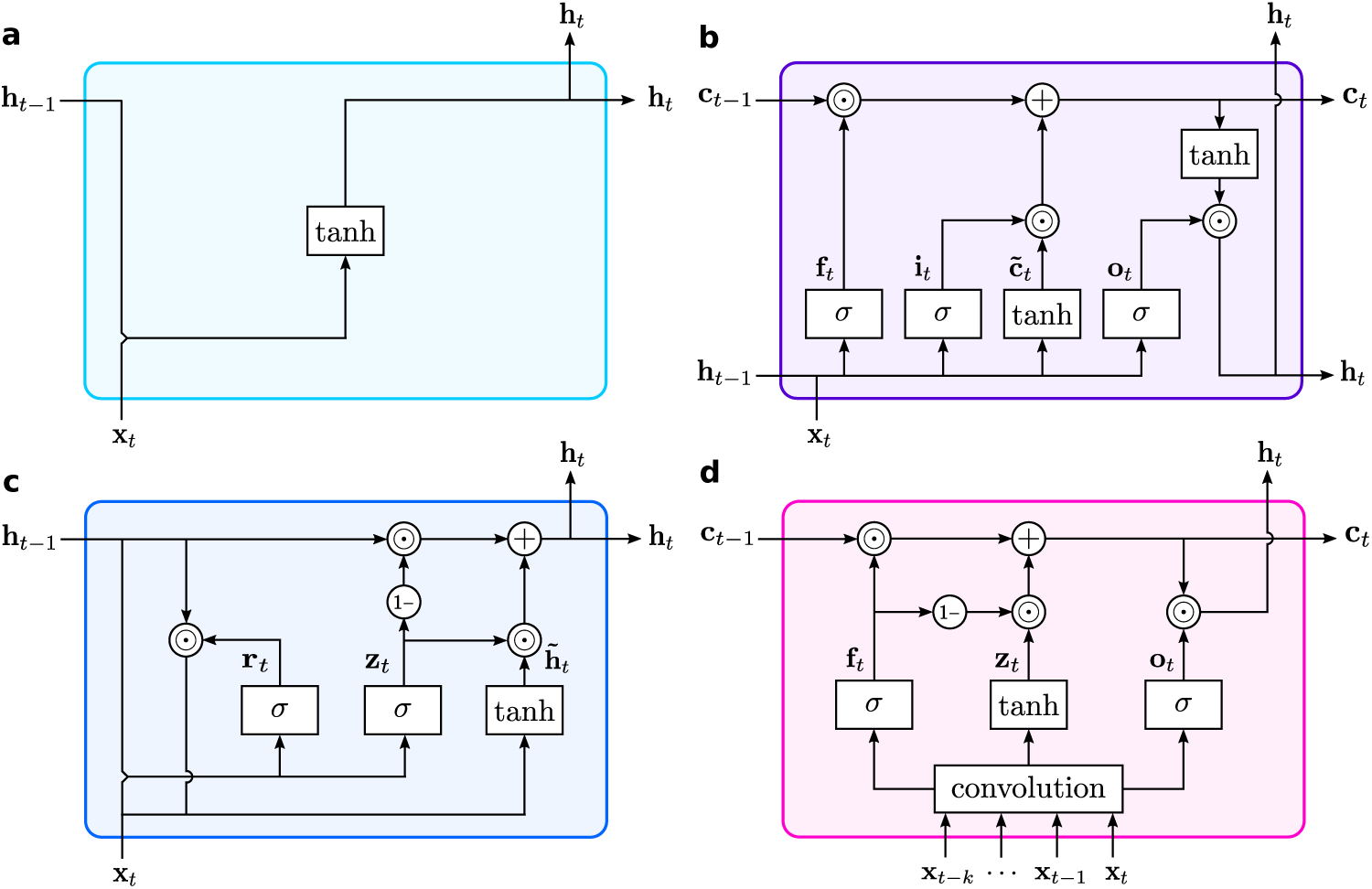
Graphical illustration of the deep learning decoders. **a**, Standard recurrent neural network (SRNN). **b**, Long short-term memory (LSTM). **c**, Gated recurrent unit (GRU). **d**, Quasi-recurrent neural network (QRNN).

**Table S1.** Hyperparameter search spaces during the optimisation of deep learning decoders.

| Hyperparameter | SRNN | GRU | LSTM | QRNN |
| --- | --- | --- | --- | --- |
| Number of units | {100, 150, …, 450} | {50, 75, …, 225} | {50, 75, …, 200} | {50, 100, …, 400} |
| Number of epochs | | {2, 3, …, 8} | | |
| Batch size |  | {32, 64, 96} |  |  |
| Dropout rate | | {0, 0.1, …, 0.5} | | |
| Learning rate | | {5, 10, …, 50} $\times 10^{-4}$ | | |

**Table S2.** Hyperparameter configurations for SUA-, MUA-, and LFP-driven deep learning decoders.

| Hyperparameter | SUA Signal |  |  |  | MUA Signal |  |  |  | LFP Signal |  |  |  |
| --- | --- | --- | --- | --- | --- | --- | --- | --- | --- | --- | --- | --- |
|  | SRNN | GRU | LSTM | QRNN | SRNN | GRU | LSTM | QRNN | SRNN | GRU | LSTM | QRNN |
| Number of units | 400 | 200 | 200 | 400 | 450 | 200 | 150 | 400 | 450 | 175 | 125 | 350 |
| Number of epochs | 8 | 3 | 6 | 4 | 5 | 4 | 6 | 4 | 7 | 3 | 5 | 2 |
| Batch size | 64 | 96 | 64 | 96 | 64 | 64 | 64 | 64 | 64 | 32 | 64 | 64 |
| Dropout rate | 0.1 | 0.4 | 0 | 0.4 | 0.1 | 0.3 | 0 | 0.4 | 0.1 | 0.3 | 0.2 | 0.1 |
| Learning rate | 0.0025 | 0.0045 | 0.002 | 0.003 | 0.0035 | 0.003 | 0.0035 | 0.005 | 0.0002 | 0.0035 | 0.004 | 0.0005 |

**Fig. S2.**
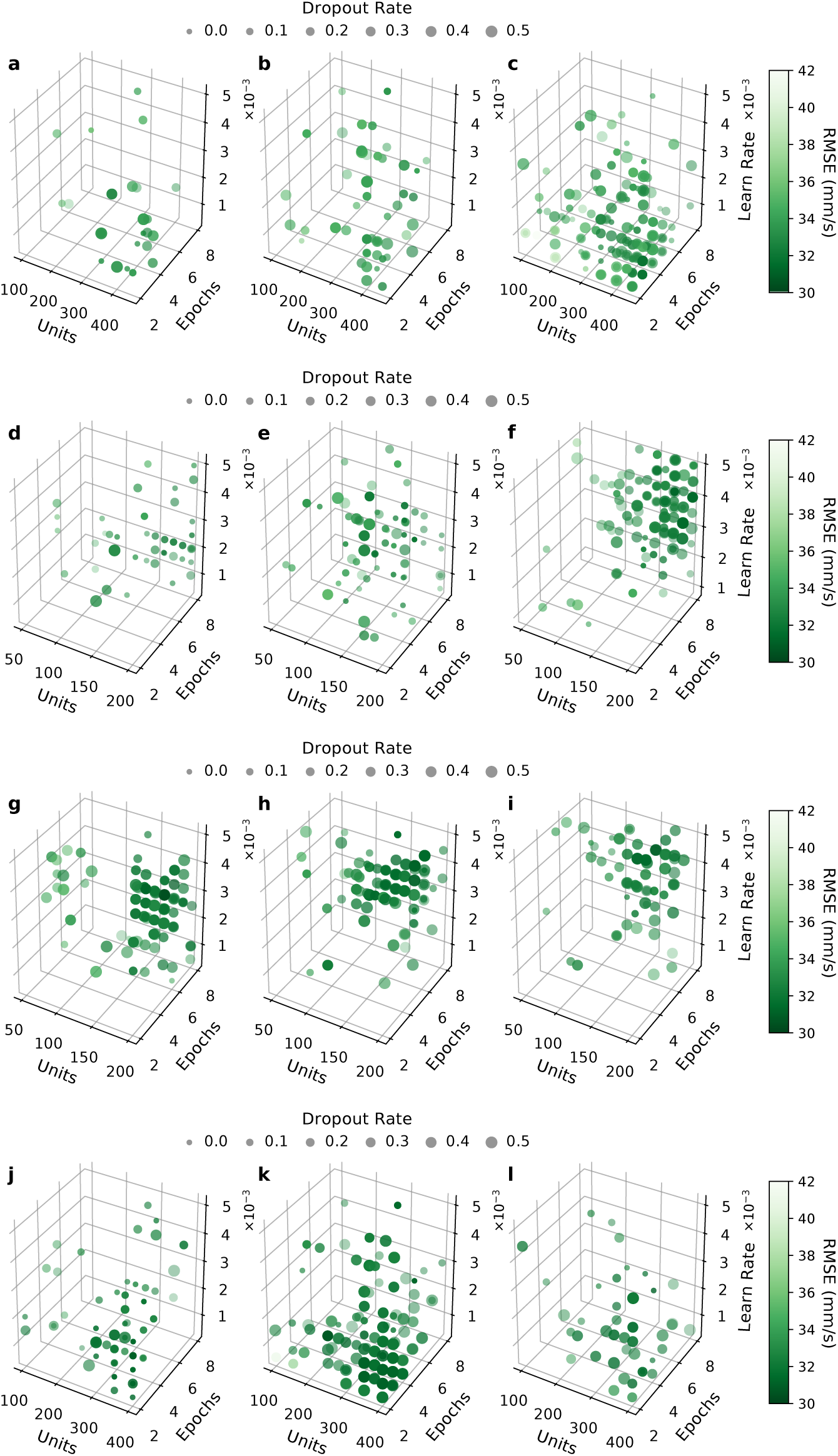
Visualisation of hyperparameter optimisation result for ESA-driven deep learning decoders. **a-c**, 3D scatter plots correspond to RNN decoder with the batch size of 32, 64, and 96, respectively. **d-f**, 3D scatter plots correspond to GRU decoder with the batch size of 32, 64, and 96, respectively. **g-i**, 3D scatter plots correspond to LSTM decoder with the batch size of 32, 64, and 96, respectively. **j-l**, 3D scatter plots correspond to QRNN decoder with the batch size of 32, 64, and 96, respectively. The *x*-axes represent the number of units; *y*-axes represent the number of epochs; *z*-axes represent the learning rate. The size of circles denotes the dropout rate while the color of circles denotes the loss value (RMSE).

**Fig. S3.**
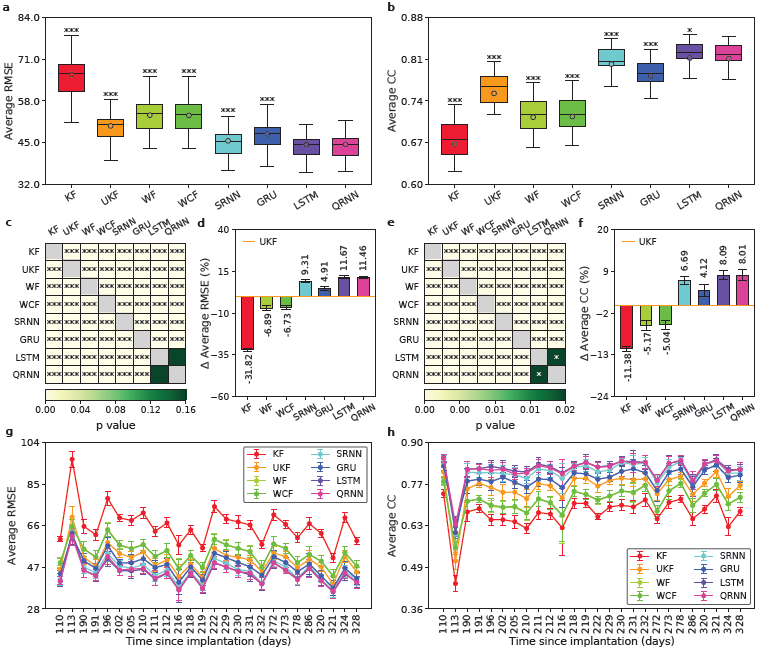
Long-term decoding performance comparison across different decoders driven by SUA signal. **a,b**, Boxplot comparison of long-term decoding performance across decoders as measured by average RMSE and CC, respectively. The horizontal lines inside the boxes denote the medians whereas the circles denote the means. Asterisks represent decoders whose decoding performance differed significantly from that of QRNN (two-tailed paired Wilcoxon signed-rank test; * *p* < 0.05, *** *p* < 0.001). **c,e**, Heatmap of statistical significant matrix of decoding performance between all possible pairs of decoders as measured by average RMSE and CC, respectively. Each cell is coloured based on the *p*-value except cells in the main diagonal (grey; excluded from statistical test calculation). Asterisks indicate whether there were statistical significant differences between the pairs (two-tailed paired Wilcoxon signed-rank test; * *p* < 0.05, *** *p* < 0.001). **d,f**, Average performance improvement/degradation (in %) of decoders with respect to that of UKF as measured by RMSE and CC metrics, respectively. Positive (negative) value indicates performance improvement (degradation). Black numeral texts represent the means whereas black error bars represent the 95% confidence intervals (*n* = 26). **g,h**, Decoding performance comparison over time across decoders in terms of average RMSE and CC, respectively. Coloured circles represent the means whereas coloured error bars represent the 95% confidence intervals (*n* = 10) for each session. The *x*-axes represent the number of days since electrode implantation. The *x*-axes’ ticks correspond to different sessions having irregular gaps between the consecutive sessions.

**Fig. S4.**
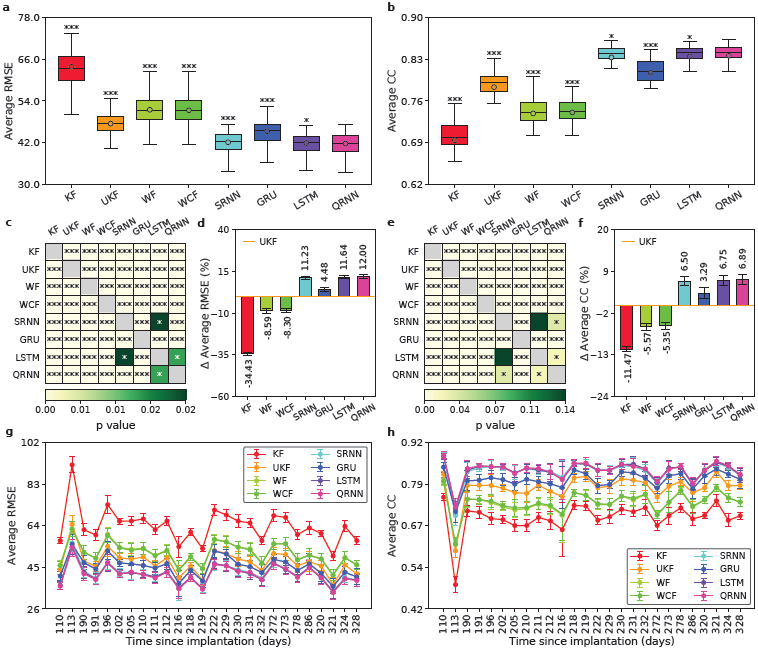
Long-term decoding performance comparison across different decoders driven by MUA signal. **a,b**, Boxplot comparison of long-term decoding performance across decoders as measured by average RMSE and CC, respectively. The horizontal lines inside the boxes denote the medians whereas the circles denote the means. Asterisks represent decoders whose decoding performance differed significantly from that of QRNN (two-tailed paired Wilcoxon signed-rank test; * *p* < 0.05, *** *p* < 0.001). **c,e**, Heatmap of statistical significant matrix of decoding performance between all possible pairs of decoders as measured by average RMSE and CC, respectively. Each cell is coloured based on the *p*-value except cells in the main diagonal (grey; excluded from statistical test calculation). Asterisks indicate whether there were statistical significant differences between the pairs (two-tailed paired Wilcoxon signed-rank test; * *p* < 0.05, *** *p* < 0.001). **d,f**, Average performance improvement/degradation (in %) of decoders with respect to that of UKF as measured by RMSE and CC metrics, respectively. Positive (negative) value indicates performance improvement (degradation). Black numeral texts represent the means whereas black error bars represent the 95% confidence intervals (*n* = 26). **g,h**, Decoding performance comparison over time across decoders in terms of average RMSE and CC, respectively. Coloured circles represent the means whereas coloured error bars represent the 95% confidence intervals (*n* = 10) for each session. The *x*-axes represent the number of days since electrode implantation. The *x*-axes’ ticks correspond to different sessions having irregular gaps between the consecutive sessions.

**Fig. S5.**
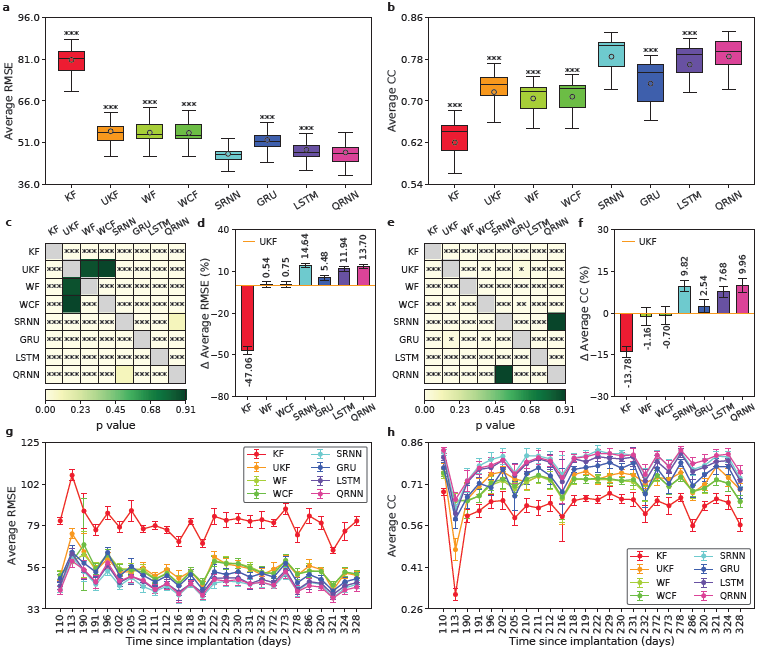
Long-term decoding performance comparison across different decoders driven by LFP signal. **a,b**, Boxplot comparison of long-term decoding performance across decoders as measured by average RMSE and CC, respectively. The horizontal lines inside the boxes denote the medians whereas the circles denote the means. Asterisks represent decoders whose decoding performance differed significantly from that of QRNN (two-tailed paired Wilcoxon signed-rank test; *** *p* < 0.001). **c,e**, Heatmap of statistical significant matrix of decoding performance between all possible pairs of decoders as measured by average RMSE and CC, respectively. Each cell is coloured based on the *p*-value except cells in the main diagonal (grey; excluded from statistical test calculation). Asterisks indicate whether there were statistical significant differences between the pairs (two-tailed paired Wilcoxon signed-rank test; * *p* < 0.05, ** *p* < 0.01, *** *p* < 0.001). **d,f**, Average performance improvement/degradation (in %) of decoders with respect to that of UKF as measured by RMSE and CC metrics, respectively. Positive (negative) value indicates performance improvement (degradation). Black numeral texts represent the means whereas black error bars represent the 95% confidence intervals (*n* = 26). **g,h**, Decoding performance comparison over time across decoders in terms of average RMSE and CC, respectively. Coloured circles represent the means whereas coloured error bars represent the 95% confidence intervals (*n* = 10) for each session. The *x*-axes represent the number of days since electrode implantation. The *x*-axes’ ticks correspond to different sessions having irregular gaps between the consecutive sessions.

**Fig. S6.**
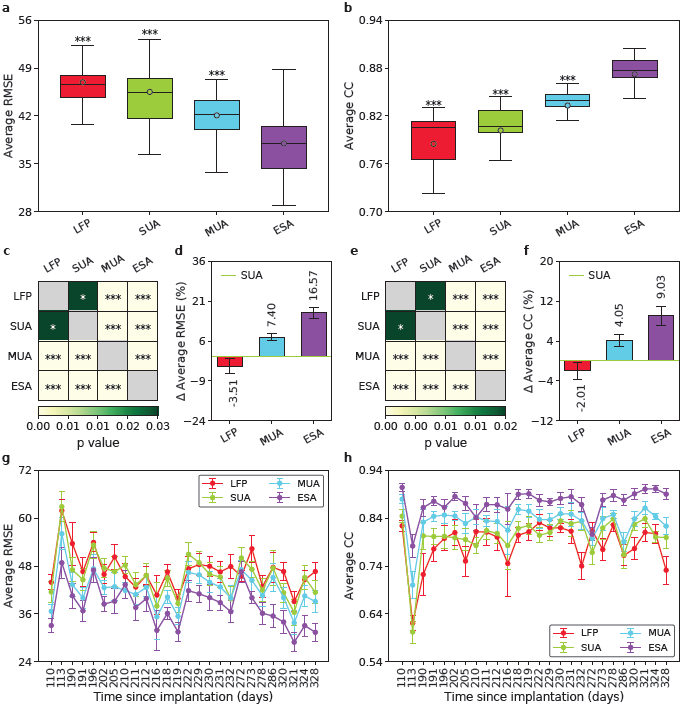
Long-term decoding performance comparison across SRNN decoders driven by different neural signals. **a,b**, Boxplot comparison of long-term decoding performance across different neural signals as measured by average RMSE and CC, respectively. The horizontal lines inside the boxes denote the medians whereas the circles denote the means. Asterisks represent neural signals whose decoding performance differed significantly from that of ESA (two-tailed paired Wilcoxon signed-rank test; *** *p* < 0.001). **c,e**, Heatmap of statistical significant matrix of decoding performance between all possible pairs of neural signals as measured by average RMSE and CC, respectively. Each cell is coloured based on the *p*-value except cells in the main diagonal (grey; excluded from statistical test calculation). Asterisks indicate whether there were statistical significant differences between the pairs (two-tailed paired Wilcoxon signed-rank test; * *p* < 0.05, *** *p* < 0.001). **d,f**, Average performance improvement/degradation (in %) of neural signals with respect to that of SUA as measured by RMSE and CC metrics, respectively. Positive (negative) value indicates performance improvement (degradation). Black numeral texts represent the means whereas black error bars represent the 95% confidence intervals (*n* = 26). **g,h**, Decoding performance comparison over time across neural signals in terms of average RMSE and CC, respectively. Coloured circles represent the means whereas coloured error bars represent the 95% confidence intervals (*n* = 10) for each session. The *x*-axes represent the number of days since electrode implantation. The *x*-axes’ ticks correspond to different sessions having irregular gaps between the consecutive sessions.

**Fig. S7.**
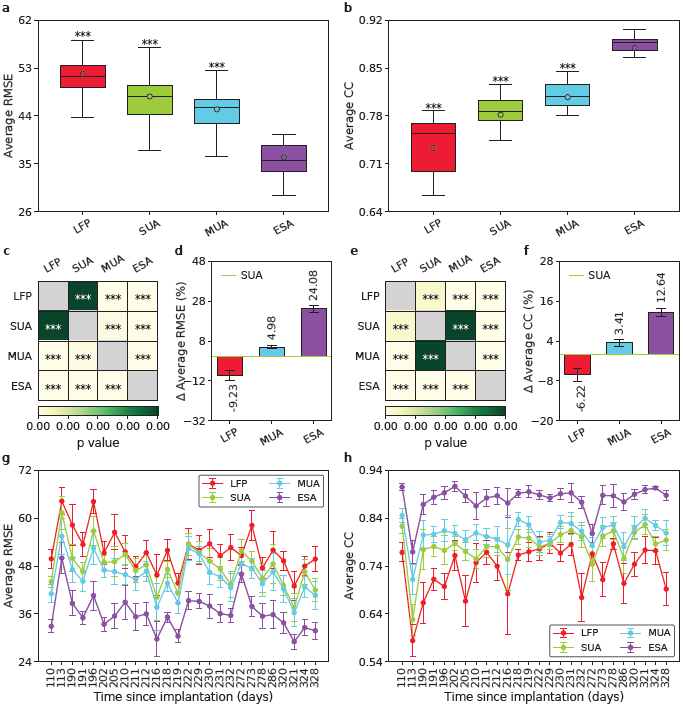
Long-term decoding performance comparison across GRU decoders driven by different neural signals. **a,b**, Boxplot comparison of long-term decoding performance across different neural signals as measured by average RMSE and CC, respectively. The horizontal lines inside the boxes denote the medians whereas the circles denote the means. Asterisks represent neural signals whose decoding performance differed significantly from that of ESA (two-tailed paired Wilcoxon signed-rank test; *** *p* < 0.001). **c,e**, Heatmap of statistical significant matrix of decoding performance between all possible pairs of neural signals as measured by average RMSE and CC, respectively. Each cell is coloured based on the *p*-value except cells in the main diagonal (grey; excluded from statistical test calculation). Asterisks indicate whether there were statistical significant differences between the pairs (two-tailed paired Wilcoxon signed-rank test; *** *p* < 0.001). **d,f**, Average performance improvement/degradation (in %) of neural signals with respect to that of SUA as measured by RMSE and CC metrics, respectively. Positive (negative) value indicates performance improvement (degradation). Black numeral texts represent the means whereas black error bars represent the 95% confidence intervals (*n* = 26). **g,h**, Decoding performance comparison over time across neural signals in terms of average RMSE and CC, respectively. Coloured circles represent the means whereas coloured error bars represent the 95% confidence intervals (*n* = 10) for each session. The *x*-axes represent the number of days since electrode implantation. The *x*-axes’ ticks correspond to different sessions having irregular gaps between the consecutive sessions.

**Fig. S8.**
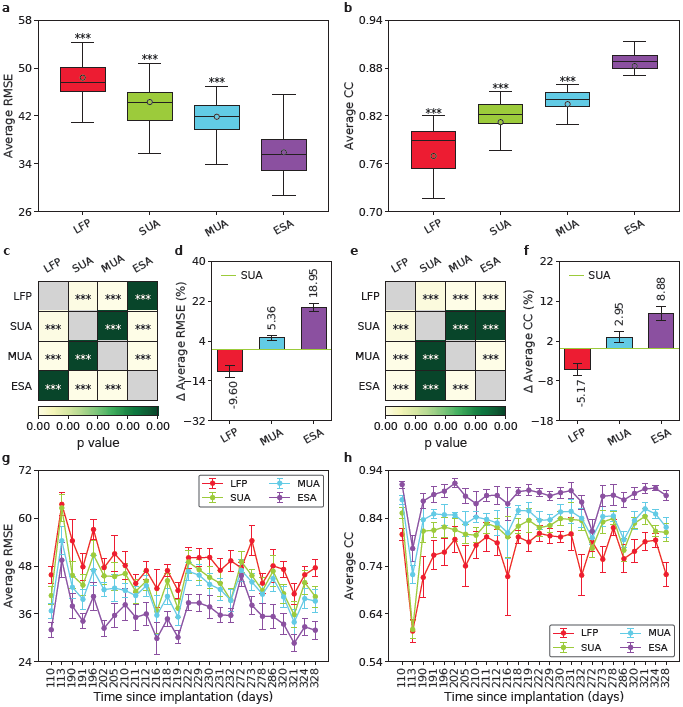
Long-term decoding performance comparison across LSTM decoders driven by different neural signals. **a,b**, Boxplot comparison of long-term decoding performance across different neural signals as measured by average RMSE and CC, respectively. The horizontal lines inside the boxes denote the medians whereas the circles denote the means. Asterisks represent neural signals whose decoding performance differed significantly from that of ESA (two-tailed paired Wilcoxon signed-rank test; *** *p* < 0.001). **c,e**, Heatmap of statistical significant matrix of decoding performance between all possible pairs of neural signals as measured by average RMSE and CC, respectively. Each cell is coloured based on the *p*-value except cells in the main diagonal (grey; excluded from statistical test calculation). Asterisks indicate whether there were statistical significant differences between the pairs (two-tailed paired Wilcoxon signed-rank test; *** *p* < 0.001). **d,f**, Average performance improvement/degradation (in %) of neural signals with respect to that of SUA as measured by RMSE and CC metrics, respectively. Positive (negative) value indicates performance improvement (degradation). Black numeral texts represent the means whereas black error bars represent the 95% confidence intervals (*n* = 26). **g,h**, Decoding performance comparison over time across neural signals in terms of average RMSE and CC, respectively. Coloured circles represent the means whereas coloured error bars represent the 95% confidence intervals (*n* = 10) for each session. The *x*-axes represent the number of days since electrode implantation. The *x*-axes’ ticks correspond to different sessions having irregular gaps between the consecutive sessions.

**Fig. S9.**
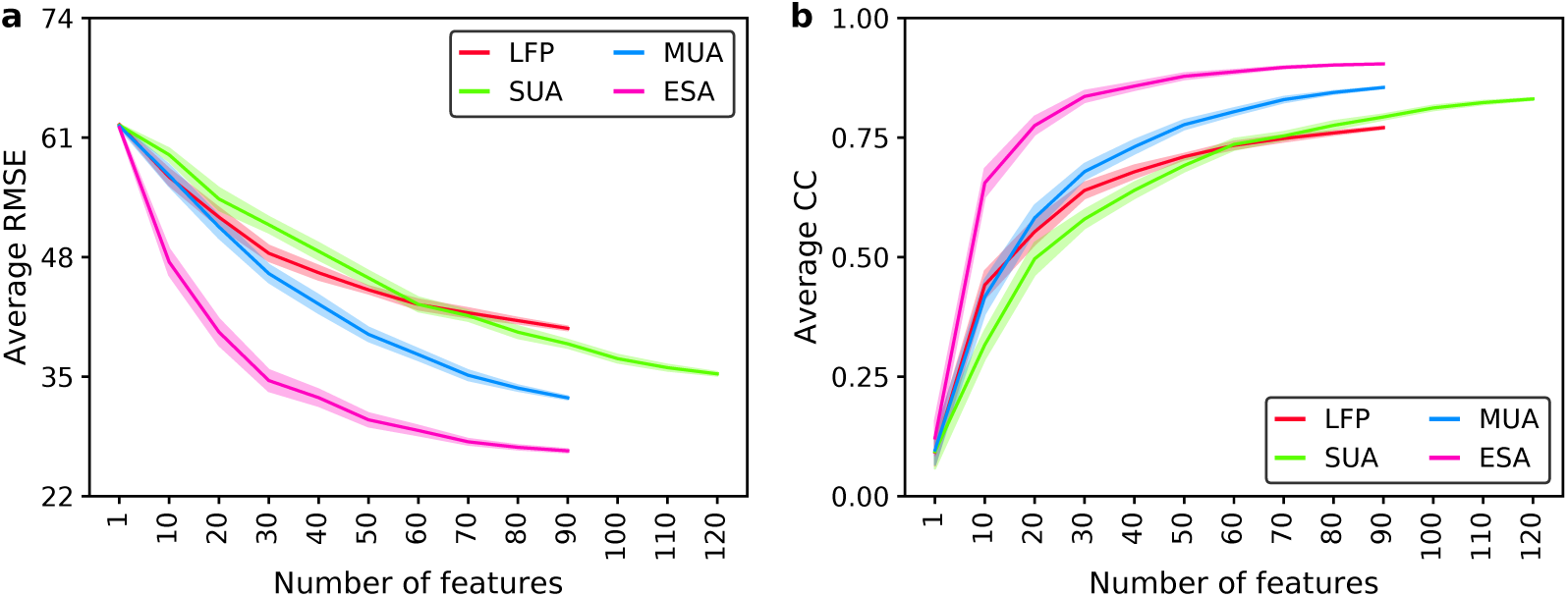
Decoding performance comparison across different types of neural signals. **a,b**, Impact of different number of features (units or channels) on the decoding comparison of QRNN decoders driven by different neural signals as measured by average RMSE and CC, respectively. The shaded areas represent the 95% confidence intervals (*n* = 30). Data are taken from the last session (I20170131_02).

